# DJ-1 deficiency in SH-SY5Y cells reveals dysregulated networks of genes and pathways involved in neuronal function and disease

**DOI:** 10.1101/2024.11.01.621572

**Authors:** Nathan Gock, Grace Kim, Tessa Morin, Amar Mahal, Julia Kostka, Timothy V. Beischlag, Frank J.S. Lee

## Abstract

Parkinson’s disease (PD) is the second most common neurodegenerative disease, affecting between 2 – 3% of the population aged 65 and older. Although the etiology of idiopathic PD is still to be elucidated, the study of heritable forms of the disease can provide new understanding into disease mechanisms. Recessively inherited loss of function mutations in the *PARK7/DJ-1* gene has been found to be causative for familial, early-onset PD. Importantly, *PARK7/DJ-1* related forms of familial PD replicate common disease phenotypes seen in idiopathic PD, including degeneration of substantia nigra dopaminergic neurons, and Parkinsonism. In this study, we evaluate the loss of function of *PARK7/DJ-1* on a human neuronal cell line, SH-SY5Y. Following ablation of the *PARK7/DJ-1* gene via CRISPR-Cas9, RNA sequencing and the DESEQ2 tool kit were utilized to filter differentially expressed genes between *PARK7/DJ-1* knockouts and control SH- SY5Y cells. 5684 genes were identified to be significantly differentially expressed. 3 genes from each of the top 10 upregulated (ATOH8, LAYN, TLX2) and downregulated (CACNA1B, CPLX2, SV2C) gene lists were selected and confirmed via RT-PCR. Differentially expressed gene lists were run through the WebGestalt functional enrichment analysis toolkit to identify enriched gene ontology (GO) terms for biological processes, cellular components, molecular function, and Kyoto Encyclopedia of Genes and Genomes (KEGG) pathways respectively. Among the top 10 significantly enriched KEGG pathways for upregulated genes were those related to neurodegenerative diseases such as Parkinson’s disease, Alzheimer’s disease, and Huntington’s disease (p-adj ≤ 0.05). Differentially expressed genes were run through the STRING database to predict protein-protein interactions (PPI). A highly significant PPI enrichment was observed (p < 1.0e-16). Our results indicate that loss of DJ-1 function in human neuronal cells leads to dysregulation of networks of connected genes and pathways that are implicated in neurodegenerative disease as well as neuronal function.

## Introduction

Parkinson’s disease (PD) is the second most common neurodegenerative disease, affecting between 2 – 3% of the population aged 65 and older (1,2). It is estimated that in the world’s 10 most populous countries, an aging population will lead to a significant increase in individuals affected by PD by 2050, causing considerable healthcare, economic, and social burden (3,4). The cellular hallmarks of PD include a loss of dopamine (DA) neurons in the substantia nigra of the midbrain, as well as the presence of aggregated α-synuclein in the form of Lewy bodies (5). Clinically, these hallmarks manifest as a progressive worsening of motor functioning symptoms, including bradykinesia, rigidity, tremor, and postural instability (6). While most PD cases are sporadic, monogenic forms of PD account for up to 10% of cases and show similar cellular pathogenesis to sporadic PD such as mitochondrial dysfunction and oxidative stress (7,8). One such monogenic, early-onset form of PD is linked to recessive mutations in the PARK7/DJ-1 gene, which codes for the 189 amino acid DJ-1 protein (9). Patients with PARK7/DJ-1 mutations demonstrate cardinal PD phenotypes, such as motor dysfunction, DA neuronal loss, and α- synuclein-positive Lewy bodies (10). The DJ-1 protein itself has been linked to a wide variety of cellular functions, including transcriptional regulation, mitochondrial function, and as a molecular chaperone (11–16). Additionally, our laboratory has previously published on the role that DJ-1 plays in DA reuptake through facilitating DA transporter (DAT) cell surface localization (17). Perhaps the most well characterized of DJ-1’s function in relation to PD is its role in oxidative stress protection (18,19). Interestingly, oxidative damage of DJ-1 is shown to be elevated in sporadic PD, further highlighting the similarities between monogenic and sporadic forms of the disease (20).

The PARK7/DJ-1 PD-related mutations are largely considered loss of function mutations as they can disrupt the normal dimerization of DJ-1 in living cells, as seen with the L166P, M26I, L10P, and P158Δ mutations (21). Furthermore, The L166P, L10P, and P158Δ mutations all lead to protein instability, and results in rapid degradation of DJ-1 (22–24). Consequently, it is important to study the downstream effects of loss of DJ-1 in biological systems. Many rodent models of DJ- 1 loss of function have been evaluated. DJ-1 knock-out (KO) rats exhibit progressive loss of DA neurons, motor dysfunction, and altered striatal DA signaling (25–27). DJ-1 KO mice, however, do not exhibit the same progressive DA degeneration or motor dysfunction. They do display increased sensitivity to oxidative stress, as well as altered DA signaling (28–30). DJ-1 KO and DJ- 1 silencing have also been evaluated in cellular models, demonstrating a wide range of impaired functioning, including with mitochondrial function, synaptic vesicle endocytosis, inflammatory responses, and most recently, ATP synthase regulation (31–35). Previous cellular models have used cultured cells from either DJ-1 KO mice tissue or from fibroblasts cultured from family members carrying DJ-1 mutations. Furthermore, much of the previous research on DJ-1 loss of function had focused on specific biological processes related to DJ-1 functioning rather than large-scale gene expression analyses. Therefore, in this study we investigate the effect of DJ-1 ablation in the human SH-SY5Y neural cell line. We utilize next-generation sequencing to explore large-scale gene expression changes between DJ-1 KO and WT human SH-SY5Y cells, and further investigate how these alterations in gene expression relate to biological processes, cellular components, and biochemical pathways.

## Materials and methods

### gRNA design and CRISPR mediated gene ablation

CRISPR/Cas9 gene editing was performed as described previously (36). In brief, sgRNAs were designed to generate PARK7/DJ-1 knock out using the online tool previously hosted at crispr.mit.edu (36) . After selecting 3 sgRNA sequences, oligos were ordered from IDTDNA and cloned into pX330-U6-Chimeric_BB-CBh-hSpCas9 (Plasmid #42230; Addgene). Plasmids were validated by Sanger sequencing (Eurofins Operon). Plasmids were transfected into SH-SY5Y cells using polyethylenimine (PEI) transfection as described previously (37,38). After low-density seeding, single clones were isolated using 5 μl of Trypsin-EDTA (Thermo Fisher Scientific) and expanded in 96-well plates. Clones were expanded and probed for DJ-1 expression using Western blots. The resulting knockout clones were named as follows: P7-1, P7-2, P7-3 and pCMV (empty control vector). KO of QPRT was further shown on RNA and protein level using real-time RT-PCR and Western Blot, respectively.

### Cell culture, transfection and colony selection

SH-SY5Y cells were cultured in DMEM (Life Technologies) supplemented with 10% FBS (Life technologies). On the day prior to transection, cells were seeded into tissue culture plates at 25- 33% confluency. On the day of transfection, media was changed to serum-free DMEM. SH-SY5Y cells were transfected with PEI (37,38). Briefly, 2 ug of pCMV-3tag8 plasmid and 6 ug of pX330 plasmid DNA containing gRNA sequences (see above) were resuspended in 500 µL of OPTI-MEM media and, in a separate tube, 20 µL of PEI reagent was resuspended in 500 µL of OPTI-MEM media. Empty pCMV-3tag8 plasmid, which contains a hygromycin resistance, was used to aid in cell selection. The DNA and PEI mixtures were combined and allowed to sit for 5 min at room temperature, prior to adding to the cells. Subsequently, cells were grown under selection pressure by the addition of hygromycin (400 µg/mL) in growth media. After 2 weeks, separate colonies were identified and were isolated. These colonies of cells were further diluted and seeded into 96 well plates to obtain single cell clones. Single cell clones were then cultured and scaled up to provide enough material for western blots to verify DJ-1 knockouts.

### Western blotting

SH-SY5Y cells were homogenized in modified RIPA buffer (50 mM Tris-HCl (pH 7.5), 150 mM NaCl, 1% NP-40, 0.5% sodium deoxycholate, 2 mM EDTA, 1 mM sodium orthovanadate, 0.1% Triton X-100) with Complete Protease Inhibitor cocktail (Roche, Indianapolis, IN) for 1 hour at 4°C. Lysates were then centrifuged at 16,100 x g for 15 minutes at 4°C and the solubilized fraction were collected and protein concentrations were determined through Bradford protein assays. Samples were subsequently diluted in SDS sample buffer, boiled for 10 min and subjected to SDS-PAGE. After solubilized proteins were subjected to electrophoresis on 10% gels (SDS–PAGE). Gels were transferred to polyvinylidenedifluoride (PVDF) membrane (Bio-Rad Laboratories, Mississauga, ON), blocked with 5% non-fat milk in TBS-T buffer (10 mM Tris-HCl, 150 mM NaCl and 0.1% Tween-20) for 1 hour at room temperature, washed three times and incubated with primary antibody, at appropriate dilutions in TBS-T, overnight at 4°C. Primary antibodies used in western blots were all purchased from Santa Cruz Biotechnology (Santa Cruz, CA) and used at a 1:200 - 1000 dilution (DJ-1, sc-27004; α-tubulin, sc-8035). Following three TBS-T washes, the blot was incubated with appropriate secondary antibody (diluted 1:10,000 in TBS-T with 0.5% milk) for 1-1.5 hours at room temperature. The blots were visualized with SuperSignal West Dura Extended Duration Substrate (Thermo Scientific, Ottawa, ON) with Dyversity image analysis system and GeneSnap image acquisition software (Syngene, Frederick, MD).

### RNA extraction, library construction and RNA sequencing

For RNA extraction, Trizol (Life Technologies, Cat. # 15596018) was used according to the manufacturer’s protocol. Total RNA samples were then frozen and sent to Canada’s Michael Smith Genome Sciences Centre, Vancouver, Canada for paired-end RNA sequencing library generation and sequencing using the HiSeq platform. RNA sequencing data was processed by the Genome Sciences Centre for quality control and for the generation BAM alignment files using BWA-MEM.

### RNA sequencing analysis

Sequencing data were uploaded to the Galaxy web platform, and we used the public server at usegalaxy.org to analyze the data (39). Prior to running the DESEQ2 analysis, gene counts for each dataset were tabulated using the featureCounts tool with the hg19 human genome assembly. Subsequently, differential gene expression analysis was performed with DESEQ2 tool in Galaxy. Resulting file was then uploaded into Excel for further analysis.

### Protein-protein interaction analysis

Using the Cytoscape bioinformatics software, a PPI network was generated using the STRING database plugin (1–3). Differentially expressed genes (DEGs) were filtered (Base mean ≥ 500), (Abs(log2(FC)) ≥ 1), (P-adj ≤ 0.05) to produce a total of 703 genes, which were run through the STRING database with a confidence score cutoff of 0.7, and organized using a force-directed layout paradigm. Force-directed layout paradigms are commonly used in generating maps for biological networks (41). A total of 539 nodes with 445 edges were returned, showing a significant PPI enrichment (p < 1.0e-16). Notably, one main cluster segregated into 176 nodes with 396 edges, which was isolated and selected for figure representation. Individual genes and smaller networks ranging from 2 – 7 nodes were excluded from representation.

### Quantitative Real-time PCR

Quantitave PCR methods were carried out as previously described (42). Briefly, about 200 ng of RNA was reverse-transcribed using Superscript II (Life Technologies). SYBR Green PCR Master Mix (Life Technologies) was used for quantification of cDNA using the StepOne Real Time PCR System (Life Technologies). Oligonucleotides used are listed in Supplemental Table SX. PCR products were run on an agarose gel for confirmation of a single band amplification at the expected size. From each experiment, the threshold levels of each amplification were adjusted to the logarithmic part of the curve for determining a Ct value. Then the Ct values were normalized with those of β-actin to obtain the relative mRNA levels. The normalized data were analyzed using a Student’s t-test, and the confidence levels were displayed as p-values.

## Results

### Transcriptomic profile of DJ-1 null SH-SY5Y cells

To investigate the normal biological role of DJ-1 in a human neuronal cell line we employed the CRISPR/Cas-9 system to create DJ-1 null SH-SY5Y cells (43). Using the Zhang laboratory gRNA tool that was previously hosted at crispr.mit.edu (36), we selected 3 gRNA sequences (PARK7-1, PARK7-2, PARK7-3) that were located at the 5’ end of the coding sequence of PARK7/DJ-1 (Figure 1A). These candidate gRNAs were also screened for specificity using BLAST (Basic Local Alignment Search Tool) algorithms to minimize the risk of off-target effects. For each gRNA, we were able to identify single cell clones for further characterization. We employed 3 separate gRNAs as part of our strategy to reduce the possibility that similar phenotypes exhibited by all 3 PARK7^-/-^ clones would be due to off-target effects. As shown in Figure 1B, the clones did not exhibit any DJ-1 protein expression as indexed by Western blot. Total RNA extracted from the PARK7^-/-^ clonal lines and from control cells were used for cDNA library creation and subsequent RNA sequencing performed by Michael Smith Genome Sciences Centre (Vancouver, Canada). The resulting transcriptomic data was utilized to examine differentially expressed genes via the DESEQ2 algorithm between PARK7^-/-^ cells and from control cells. As shown in Figure 2A, there are significant levels of differential expressed genes (DEG) that segregate between the PARK7^-/-^ cells and control SH-SY5Y cells. Shown in Figure 2A are the top 500 DEGs that were identified after thresholding for genes with P values greater than 0.01 and with base mean expression greater than 500. Furthermore, volcano plots of the DEG analysis highlights several genes that show increased or decreased expression in PARK7^-/-^ cells (Figure 2B). To verify the DEG analysis, we performed real-time quantitative PCR (RT-qPCR) on 3 of the top 10 genes identified to be upregulated (Figure 3A) and downregulated (Figure 3B) in the PARK7^-/-^ cells. All 3 of the upregulated genes chosen (ATHO8, LAYN, TLX2) showed significant upregulation in gene expression in all 3 PARK7^-/-^ cells (PARK7-1, PARK7-2, PARK7-3; Figure 3A). Similarly, all 3 downregulated genes chosen (CACNA1B, CPLX2, SVC2) showed decreased mRNA levels in all 3 PARK7^-/-^ cells (PARK7-1, PARK7-2, PARK7-3) as quantified by RT-qPCR (Figure 3B).

**Figure 1.**
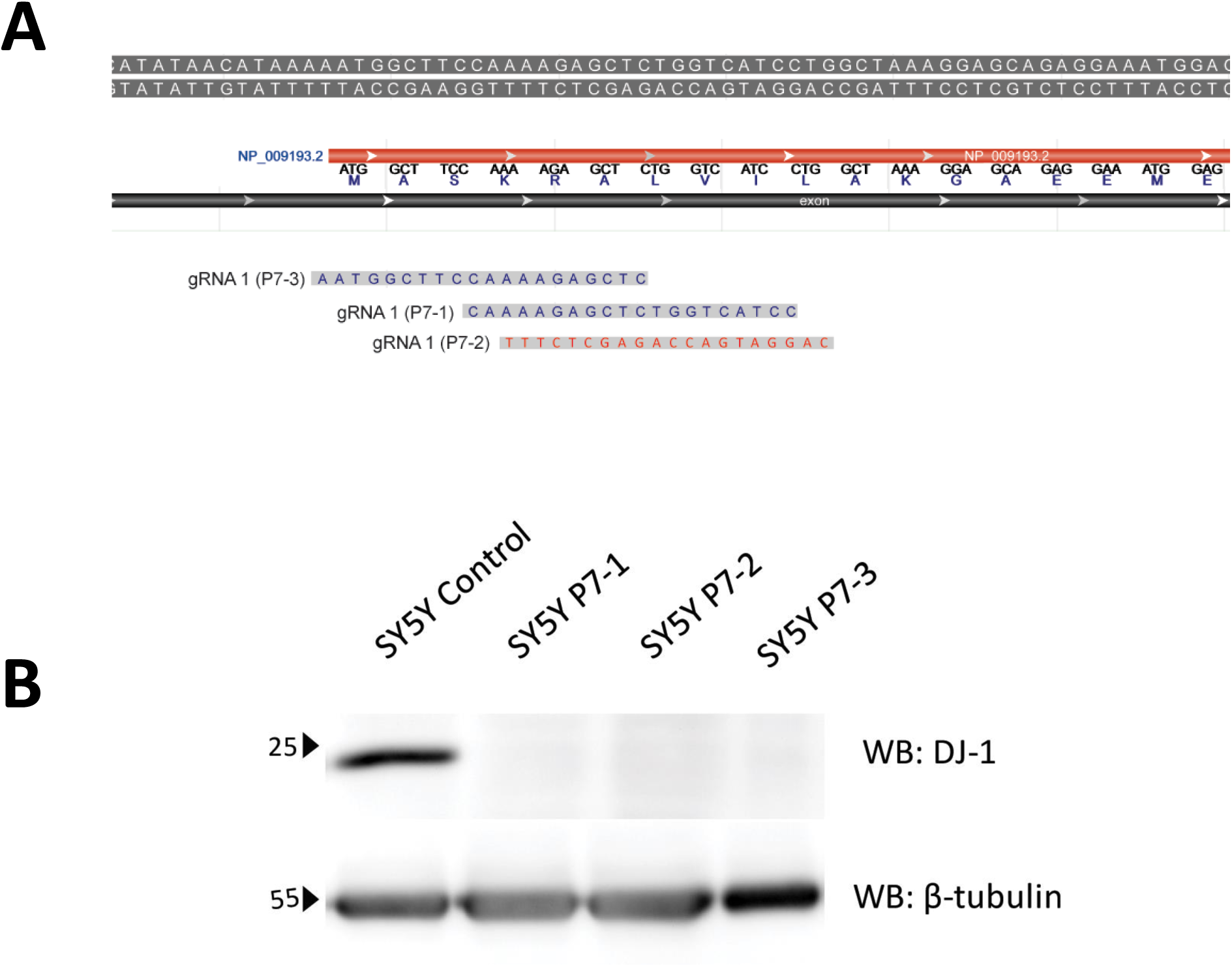
CRISPR-Cas9 mediated knockout of the PARK7/DJ-1 gene in SH-SY5Y cells. (a) Top panel illustrates the sequences that were targeted in the creation of the sgRNA used by the CRISPR-Cas9 system to disrupt the PARK7/DJ-1 gene. Blue sequences (gRNA1, gRNA3) were targeting sense strand of the PARK7/DJ-1 gene while the red sequence (gRNA2) represent utilization of the antisense sequence. Synthetic oligos were used to subclone gRNA sequences into the pX330 gRNA-Cas9 plasmid. These plasmids were then transfected into SH-SY5Y cells to disrupt the PARK7/DJ-1 gene. (b) Bottom panel are western blots for DJ-1 and b-tubulin to examine the expression if SH-SY5Y control cells and in DJ-1 KO cells (P7-1, P7-2 and P7-3).

**Figure 2.**
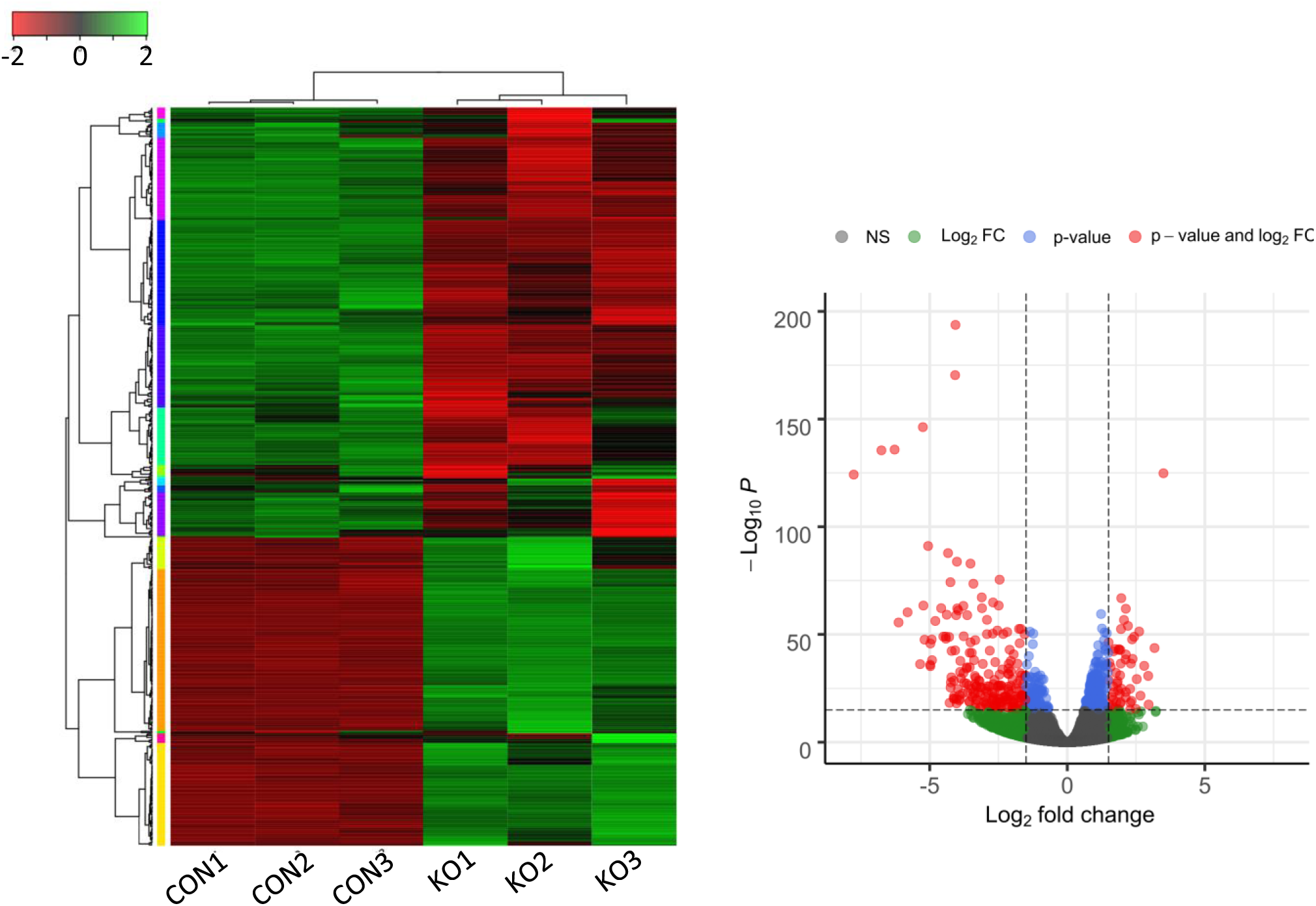
RNA sequencing and DESEQ2 analysis between control SH-SY5Y cells and PARK7/DJ-1 null cells. (A) Heatmap and clustering analysis of top 500 differentially expressed genes (DEG) from the RNA sequencing data. Cutoff used to identify DEG included P value < 0.01 and a base mean expression > 500. (B) Volcano plot of DEG between control and PARK7/DJ-1 null SH-SY5Y cells. Log2 fold change was plotted against the DESEQ2- generated p-value (-log base 10) using a P < 0.05 cutoff and log2-fold change > |1|.

**Figure 3.**
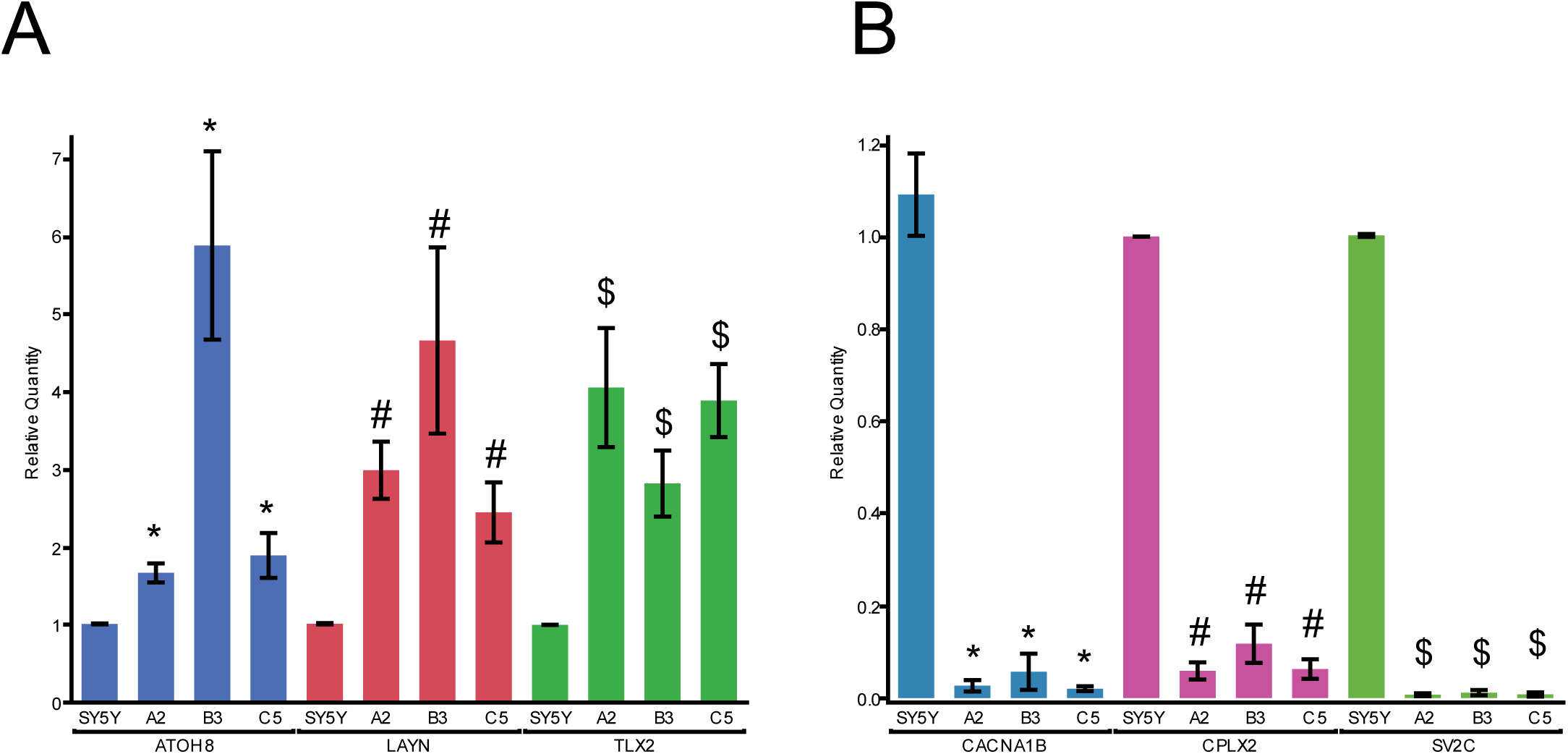
Real time PCR confirmation of RNA sequencing analysis. We chose 3 of the top 10 genes identified to be upregulated (A) and downregulated (B) in the PARK7/DJ-1 null cells. Results showed significant differences between DJ-1 KO null groups (A2, B3, C5) vs SH-SY5Y wildtype cells using nonparametric Kruskal-Wallis tests. *, #, $; P < 0.05 compared to wildtype (WT) control for respective gene indicated.

### Functional enrichment analysis on differentially expressed genes

To investigate the functional associations of the common DEGs, we performed separate GO and KEGG analyses for upregulated and downregulated genes using Over-Representation Analysis (ORA) method in WebGestalt (http://webgestalt.org) (44–46). The top 10 terms enriched in the upregulated genes are listed in Figure 4 while the top 10 terms enriched in downregulated genes are shown in Figure 5. The top GO terms (i.e., Biological Processes (BP), Cellular Components (CC), Molecular Function (MF)) in the upregulated gene list includes protein targeting (GO:0006605, BP), mitochondrial matrix (GO:0005759, CC), and cell adhesion molecule binding (GO:00050839, MF). For the downregulated gene list, the top terms for BP, CC, and MF were mRNA processing (GO:0006397, BP), axon part (GO:0033267, CC), protein serine/threonine kinase activity (GO:0004674, MF). We also explored KEGG pathways with the top results being related to Ribosome (hsa0310; upregulated gene list) and Phosphatidylinositol signaling system (hsa04070; downregulated gene list). Notably, among the upregulated gene list in the KEGG pathways analysis several neurological disorders are listed including Huntington’s, Alzheimer’s and Parkinson’s disease.

**Figure 4.**
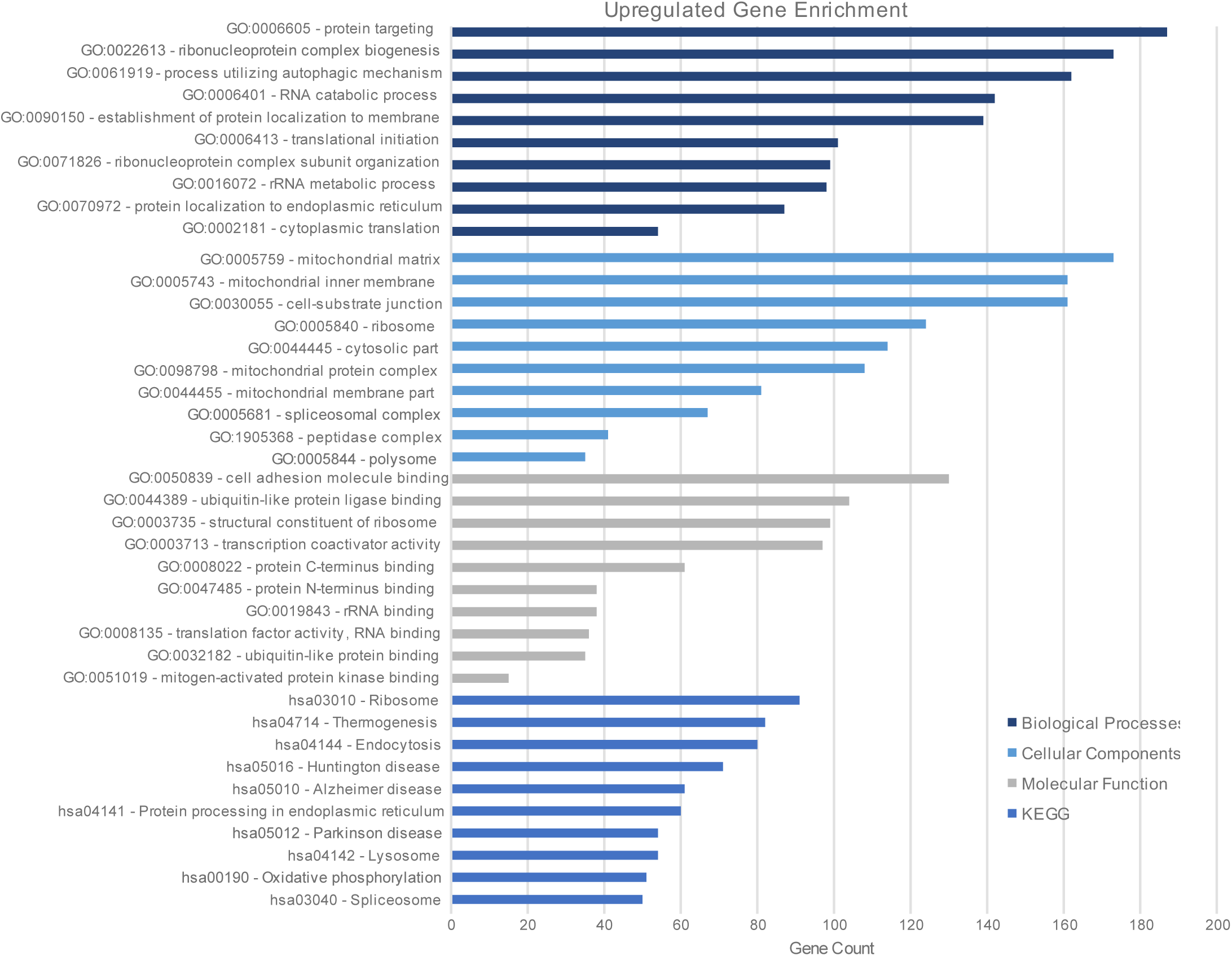
Top GO terms enriched for upregulated genes chosen by significance. RNA-Seq terms were filtered through the DESeq2 package. Selection criteria were chosen to exclude genes without power (Base mean ≥ 250), to include upregulated genes (log2(FC) > 0), and to include only significant differentially expressed genes. (P-adj ≤ 0.05). Remaining genes were then run through the WebGestalt functional enrichment analysis toolkit for GO terms in the chosen functional databases: Biological Processes, Cellular Components, Molecular Function, and KEGG pathways. Significance threshold was set at FDR < 0.05. Reflected above are the top 10 terms for each category.

**Figure 5.**
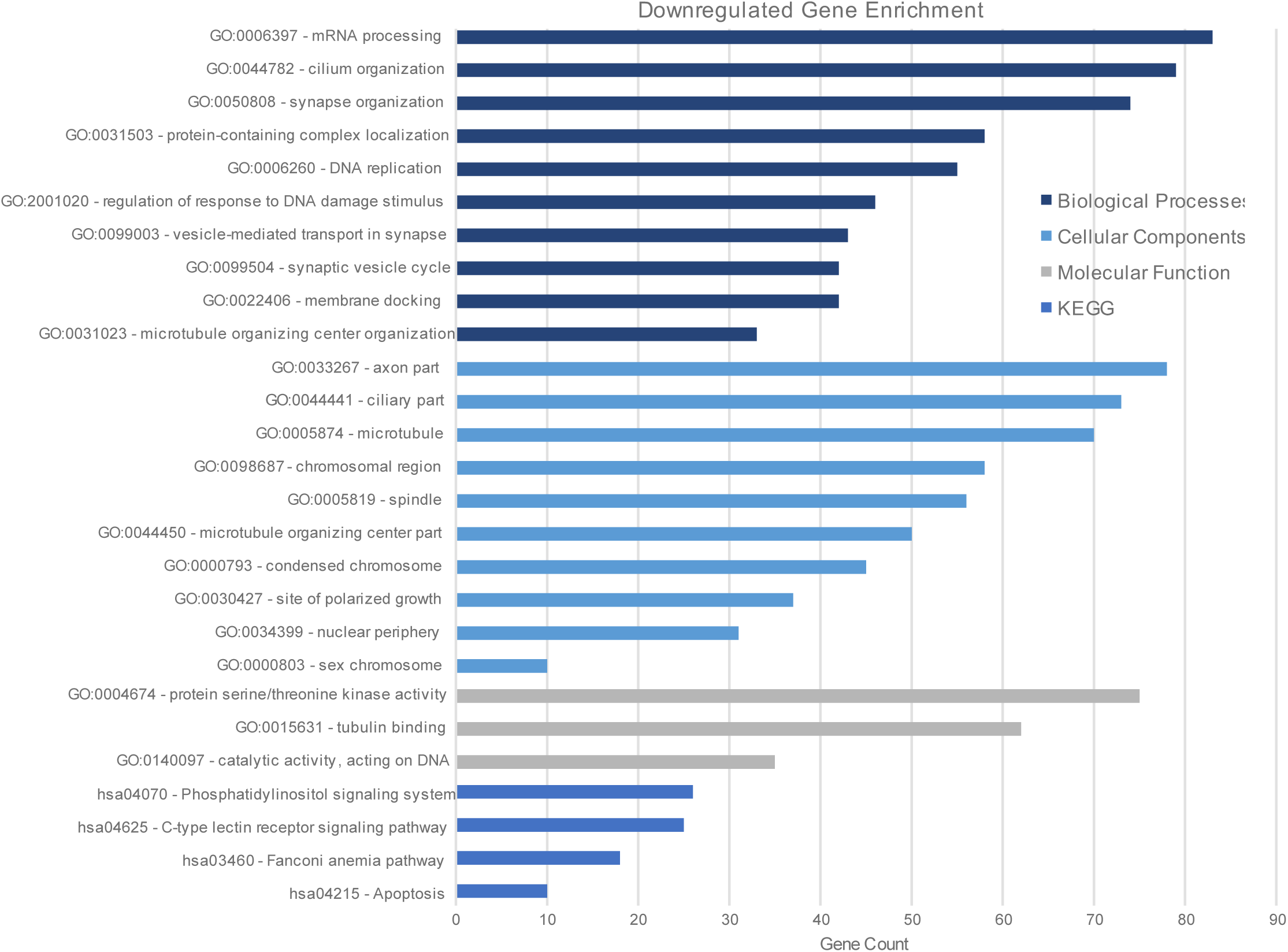
Top GO terms enriched for downregulated genes chosen by significance. RNA-Seq terms were filtered through the DESeq2 package. Selection criteria were chosen to exclude genes without power (Base mean ≥ 250), to include downregulated genes (log2(FC) < 0), and to include only significant differentially expressed genes. (P-adj ≤ 0.05). Remaining genes were then run through the WebGestalt functional enrichment analysis toolkit for GO terms in the chosen functional databases: Biological Processes, Cellular Components, Molecular Function, and KEGG pathways. Significance threshold was set at FDR < 0.05. Reflected above are the top 10 terms for Biological Processes and Cellular Components and the top terms for Molecular Function and KEGG pathways.

### PPI network and pathway deviation score analyses

In addition to functional enrichment analysis, we used our filtered DEG list to analyze protein-protein interactions (PPI) between DEG-encoded proteins using the STRING online database and the subsequent PPI network was visualized using CytoScape software. The largest continuous cluster in the PPI network had 617 nodes and 356 edges exhibiting significant PPI enrichment (P< 1.0e-16, Figure 6). Furthermore, Markov Clustering (MCL) algorithm was used to identify clusters within the network, with the top 5 clusters shown in Figure 7. Subsequent functional enrichment analysis of each cluster with CytoScape identified that the top 10 terms in cluster 1 are enriched with genes related to protein translation (Table 1). Cluster 2 is related to presynaptic neurotransmitter release pathways while cluster 3 is related to the neuron, with several terms related to action potential generation. The top terms in cluster 4 are related to apoptosis, while cluster 5 are enriched with terms related morphogenesis and development pathway related genes.

**Figure 6.**
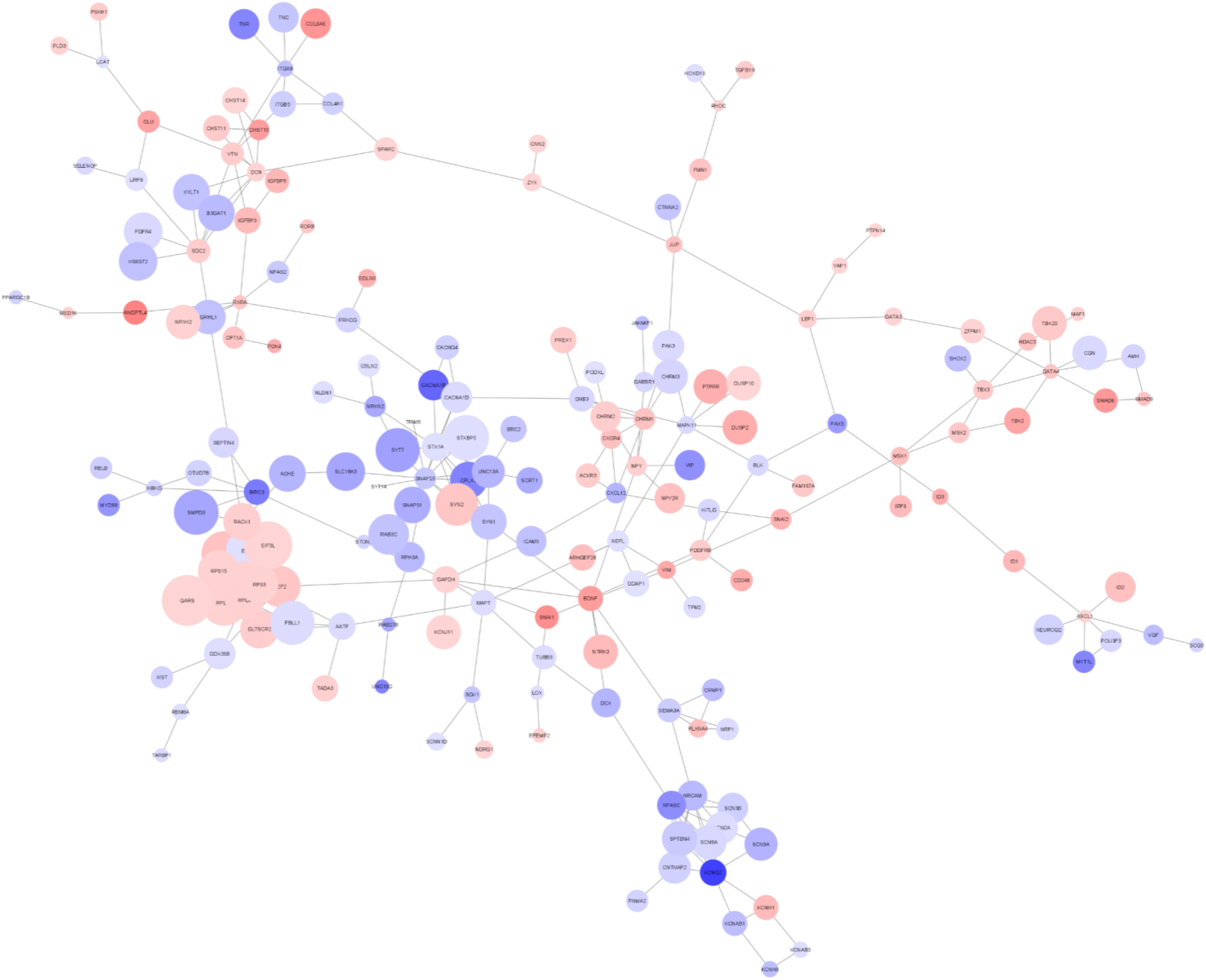
Protein-protein interaction (PPI) network of DEGs. DEGs were filtered (Base mean ≥ 250, log2(FC) ≥ |1|, P-adj ≤ 0.05) to produce 703 genes, and run through the STRING database with a confidence score cutoff of 0.7. String returned 617 nodes with 356 edges, showing a significant PPI enrichment (p < 1.0e-16). Red and blue circles represent upregulated and downregulated genes, respectively. Circles were sized according to the neighbour connectivity. Individual genes and smaller networks ranging from 2 – 7 nodes were excluded, leaving one large cluster remaining of 188 nodes, 298 edges.

**Figure 7.**
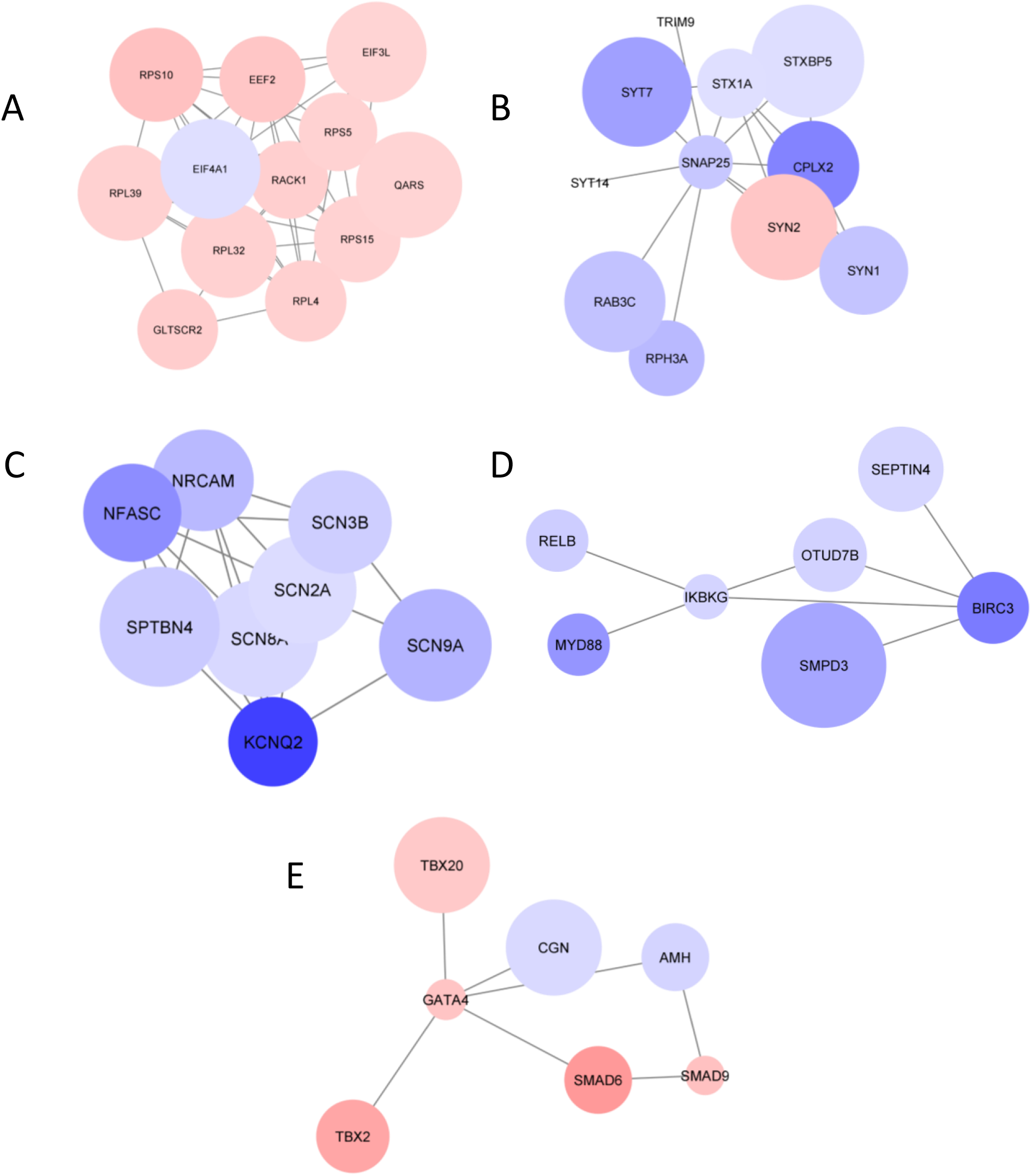
Top 5 MCL clusters identified in the protein-protein interaction (PPI) network from DEGs. In the PPI network identified through the Cytoscape STRING database, the MCL clustering algorithm (with granularity parameter /inflation value of 2.5) was used to identify clusters. Shown are the top 5 clusters (cluster 1 (A), cluster 2 (B), cluster 3 (C), cluster 4 (D), cluster 5 (E)), with red and blue circles representing upregulated and downregulated genes, respectively. Circles were sized according to the neighbour connectivity.

## Discussion

While mutations in the PARK7/DJ-1 gene have been known to lead to autosomal recessive forms of PD, there are several studies that show DJ-1 may be related to other conditions including immune and inflammatory diseases, and cancer (47–53). This is likely from DJ-1 having a multitude of cellular activities including activation of the PI3K/Akt pathway (54–58), induction of antioxidant response via Nrf2 stabilization (59–61), ERK1/2 pathway activation (62–65), ASK1 pathway inhibition (66–69) and p53 (13,70–73). To provide insight into the physiological pathways that DJ-1 is involved in, we used the CRISPR/Cas-9 system to successfully generate three PARK7-/- clonal lines that lacked DJ-1 protein expression (Figure 1). These clonal lines were compared with control SH-SY5Y cells by RNA sequencing and differential gene expression analysis using DESEQ2 (Figure 2). We found that DJ-1 deficiency resulted in significant changes in the expression of thousands of genes, implicating DJ-1’s importance in human neuronal cell transcriptome.

The results of this study show that DEGs that are common between the three PARK7^-/-^ clonal lines but differentially expressed from control SH-SY5Y cells have functional associations with various biological processes, cellular components, molecular functions, and pathways. By performing GO and KEGG analyses, we identified the top 10 terms and pathways that were enriched in our upregulated and downregulated gene lists. Upregulated genes were mainly involved in protein targeting, mitochondrial matrix, cell adhesion molecule binding, and ribosomal function. Interestingly, DJ-1 itself has been shown to have altered protein localization to mitochondria in response to oxidative stress (14,74). This may facilitate the interaction of DJ-1 with PINK1 and Parkin to facilitate mitophagy in response to oxidative damage to the mitochondria (15,75,76). One potential candidate that may facilitate the interaction between these 3 proteins is mortalin. Mortalin (or HSPA9) is a chaperone protein that has been shown to interact with DJ-1 and promote DJ-1 localization into mitochondria (77–79). Mortalin has also been shown to interact with parkin and PINK1 (80,81). Moreover, there is increasing evidence for the juxtaposition of ribosomes and mitochondria (82), which potentially could be impacted with the loss of DJ-1. We have also found that upregulated genes were enriched in pathways related to neurodegenerative diseases, such as Huntington’s disease, Alzheimer’s disease, and Parkinson’s disease (Figure 4).

This indicates that DJ-1 deficiency may alter the expression of genes that are implicated in neuronal dysfunction and degeneration and that DJ-1 plays a crucial role in maintaining neuronal function and the prevention of neurodegeneration.

On the other hand, we found that the downregulated genes were mainly involved in mRNA processing, axon components, protein serine/threonine kinase activity, and phosphatidylinositol signaling system (Figure 5). Interestingly, Bonafati et al, who discovered mutations in DJ-1 are linked to a recessive familial form of PD, speculated that these mutations could impair DJ-1 interaction with RNA binding protein complexes that impact transcription and post-transcriptional processes (9). DJ-1 itself has been shown to bind RNA in an oxidative dependent manner to regulate the mRNA translation of targets related to oxidative stress response and apoptosis (83). In addition, Repici *et al.* showed increased DJ-1 localization in stress granules after stress induction (84), whereby DJ-1 has been found to interact with mRNA of translation factors.

Furthermore, Nrf2 stabilization by DJ-1 (59) has been implicated in axon growth by regulating microtubule dynamics (85,86). DJ-1 has also been shown to impact protein phosphorylation pathways via interactions with Akt, GSKβ, Erk1/2 to name a few (54–57,57,58,62–65,87). Interestingly, these proteins are also implicated in axon growth and maintenance (88–94). Taken together, these findings suggest that DJ-1 deficiency may affect mRNA splicing, axonal structure and function, protein phosphorylation, and intracellular signaling in human neuronal cells. These processes are essential for neuronal development, plasticity, and communication.

In addition to functional enrichment analysis, we also performed protein-protein interaction analysis to examine the interactions between the DEG-encoded proteins. We used the STRING online database to construct a PPI network and visualized it using the CytoScape software (95). We found that the PPI network had a significant enrichment of interactions and formed a large cluster of connected nodes. We also used the MCL algorithm to identify sub-clusters within the network and performed functional enrichment analysis for each sub-cluster. We found that the sub-clusters had distinct functional themes that were related to neuronal function and disease. For example, cluster 1 was enriched with genes (eg. SV2C, CPLX2, SYT1, SNAP25, STXBP1, SLC18A2) encoding proteins related to presynaptic neurotransmitter release pathways, including synaptic vesicle trafficking, docking, fusion, and recycling. Cluster 2 was enriched with genes related to protein translation (RPL10A, RPL13A, RPL18A, RPL23A, RPL27A, RPL36A) that are part of the ribosomal subunits that catalyze protein synthesis. Cluster 3 was enriched with genes related to morphogenesis and development pathways (ATOH8, LAYN, TLX2, FGF13, FGF14, FGF17) that have roles in neuronal differentiation, migration, survival, and patterning. Cluster 4 was enriched with genes related to action potential generation pathways (CACNA1B, CACNA1D, CACNA1E, CACNA1G, CACNA1H, CACNA1I), which encode voltage-gated calcium channels that mediate calcium influx and trigger neurotransmitter release.

While our study provides some interesting insights into DJ-1’s physiological role, there are some limitations to our study. Our study used only one human neuronal cell line, SH-SY5Y, which may not fully represent the diversity and complexity of human neurons in vivo. Therefore, the results may not be generalizable to other neuronal cell types or to the whole brain. Although our study used three different gRNAs and screened for specificity, it did not perform whole-genome sequencing or validation of the clonal lines to confirm the absence of off-target effects or mutations. Therefore, we cannot eliminate the possibility of off-target effects or unintended mutations in other genes. Finally, while our study used functional enrichment analysis and protein-protein interaction analysis to infer the functional associations of the DEGs, this may not capture the causal relationships or the dynamic interactions of the genes and pathways. Despite these caveats, we conclude that our results demonstrate that DJ-1 deficiency leads to changes in gene expression and protein-protein interaction networks that are associated with various neuronal processes and pathways. Our results also reveal novel functional associations of DJ-1 with genes and pathways that are implicated in neurodegeneration and provides new insights into the molecular mechanisms of DJ-1 function in human neuronal cells and its potential implications for neurodegenerative disorders.

---

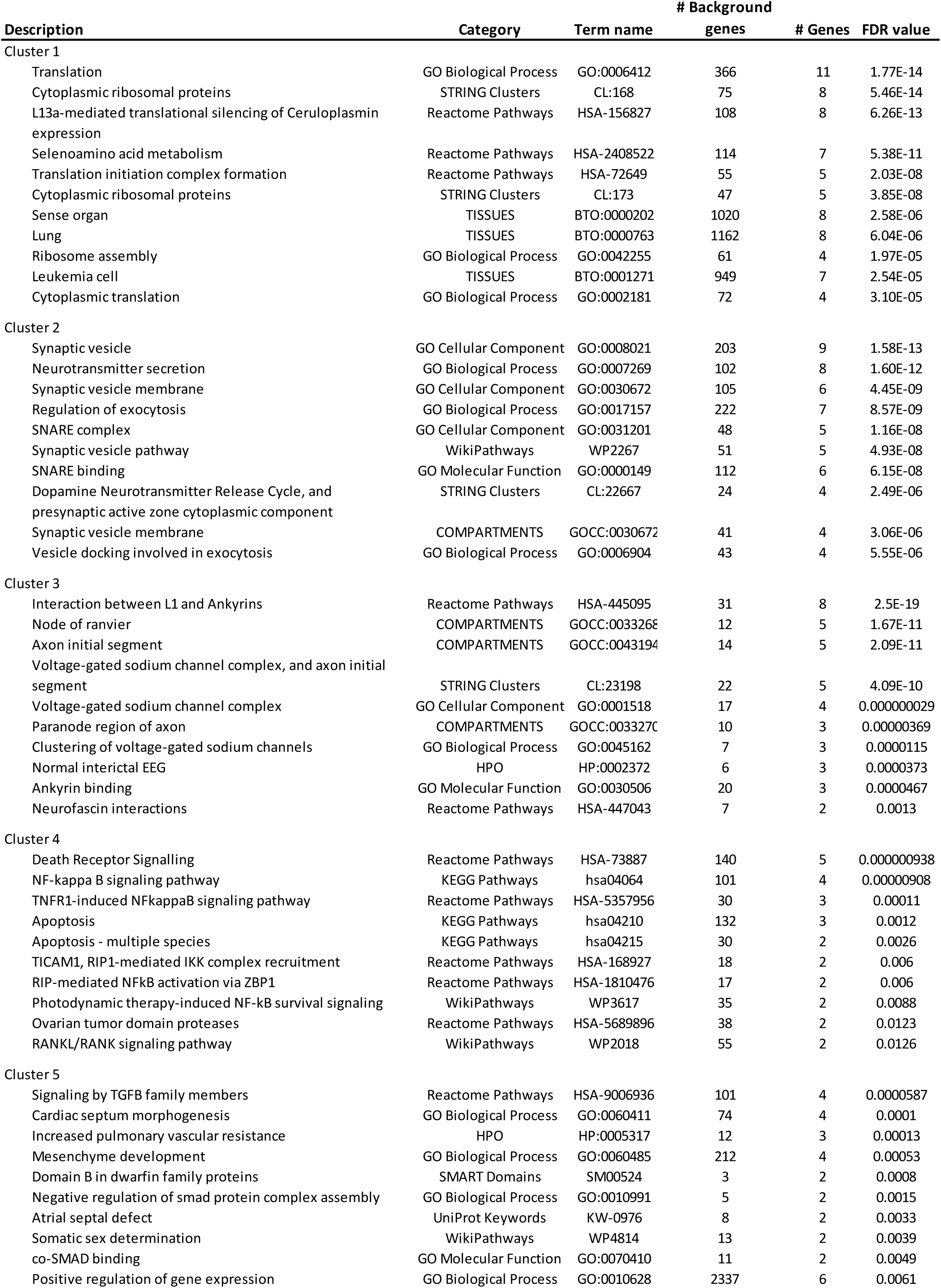

